# PLiCat: Decoding protein-lipid interactions by large language model

**DOI:** 10.1101/2025.09.09.675043

**Authors:** Feitong Dong, Jingrou Wu

## Abstract

**Motivation:** Protein-lipid interactions are essential for many cellular processes, as proteins associate with diverse lipid molecules to exert distinct functions. However, existing approaches are limited in discriminating among lipid categories in these interactions. Recent advances in protein language models enable discovery of novel sequence insights.

**Results:** We introduce PLiCat (Protein–Lipid interaction Categorization tool), a predictive framework designed to classify lipid categories interacting with proteins. By integrating ESMC and BERT models into a hybrid architecture, PLiCat achieves accurate and interpretable predictions to distinguish lipid-binding signatures across eight major lipid categories. Furthermore, we investigated the potential of PLiCat to identify lipid-binding sites and assess the impact of pathogenic mutations on lipid-binding events. Collectively, PLiCat provides a powerful framework for elucidating the lipid-binding codes, offering new opportunities for exploring lipid-binding specificity and guiding rational protein design.

**Availability and implementation:** The PLiCat source code and processed datasets are available at https://github.com/Noora68/PLiCat.

## Introduction

Lipids are a class of amphipathic biomolecules, playing crucial and diverse roles in numerous cellular processes (Harayama and Riezman 2018). In addition to the well-known function of lipids in acting as a structural component of biological membranes and reservoirs for energy storage, lipids are increasingly recognized for their involvement in signal transduction, immune homeostasis, and epigenetic regulation (Yoon *et al*. 2021). These functions are largely mediated through interactions with proteins, which participate in a wide range of physiological and pathological events (González-Becerra *et al*. 2019).

Many proteins exhibit selectivity to associate with specific lipid categories, and such lipid-binding specificity is a critical determinant of protein structure, function, and regulation (Ernst *et al*. 2010). According to the lipid classification system of LIPID MAPS, lipids can be broadly classified into eight major categories: Fatty Acyls (FA), Glycerolipids (GL), Glycerophospholipids (GP), Sphingolipids (SP), Sterol Lipids (ST), Prenol Lipids (PR), Saccharolipids (SL), Polyketides (PK) (**Supplementary** Figure 1A) (Conroy *et al*. 2024, Sud *et al*. 2007). Each category possesses distinct molecular structures and physiological functions. Specific protein-lipid interactions can modulate protein structure and function in a variety of ways (Fahy *et al*. 2011). Therefore, understanding the patterns of protein-lipid interaction is essential for unraveling the intricate biological processes.

Traditional approaches for the identification of protein-lipid interactions are highly dependent on biochemical, biophysical, and structural methods, which are time-consuming and laborious (Sych *et al*. 2022). Moreover, the essential information may have been overlooked and lost during the experimental processes. Alternatively, computational approaches for the prediction of lipid binding have been developed mainly based on the known lipid-binding domain or the physicochemical properties of amino acids, which limits our ability to discover novel protein-lipid interactions (Stahelin 2009). Due to the flexibility and complexity of protein-lipid interacting scenarios, it has been challenging to predict the preference of a certain protein for different lipid categories. Therefore, developing a predictive tool to identify the protein-associated lipid species is valuable in both biological research and protein design.

More recently, the notable breakthrough of deep-learning technologies and large language models (LLMs) has shown promising capabilities in understanding and processing biological data, providing new insights for deciphering biological languages and possibilities for capturing sequence–function relationships directly from raw amino acid sequences (Heinzinger and Rost 2025, Lin *et al*. 2023, Vaswani *et al*. 2017). For example, protein language models (PLMs), including Evolutionary Scale Modeling (ESM), conceptualize protein sequences as a language, have been pretrained on large-scale protein sequence datasets (Hayes *et al*. 2025, Lin *et al*. 2023). The emerging PLM models are advancing the field and showing immense potential in various downstream tasks, such as the prediction of protein structure, antigen-antibody binding, and protein subcellular localization (Hayes *et al*. 2025, Kilgore *et al*. 2025, Lin *et al*. 2023, Ødum *et al*. 2024, Teufel *et al*. 2022, Zhang *et al*. 2025). However, the performance of PLMs on the prediction of protein-lipid interactions has not been explored. An intriguing question is whether the PLM could discover the previously unrecognized protein code in the amino acid sequences that govern lipid binding.

To address the aforementioned issues, we present a PLM-based model, termed the Protein-Lipid interaction Categorization tool (PLiCat), leveraging the ESM Cambrian (ESMC) model (Hayes *et al*. 2025) and the Bidirectional Encoder Representation from Transformers (BERT) model (Devlin *et al*. 2019), trained on a comprehensive lipid-binding protein dataset. Particularly, our model achieves superior performance in the classification task of distinguishing different lipid-binding categories. Furthermore, it provides residue-level interpretability by identifying lipid-binding sites through attribution analysis. Finally, we evaluated the effects of pathogenic mutations that may alter protein-lipid interactions. Overall, by using protein sequences as the sole input, PLiCat offers an efficient and accurate approach to predict the lipid-binding categories, aids in decoding the sequence determinants underlying lipid recognition, and facilitates the understanding of lipid-associated protein functions and their therapeutic implications.

## Results

### PLiCat is a novel architecture tailored for predicting protein–lipid interactions

To investigate the codes hidden in protein sequences that are responsible for lipid binding, we set out to develop PLiCat, a predictive tool to distinguish sequence features of proteins binding to different lipid categories by using protein sequence as the only input.

As illustrated in Figure 1, we constructed PLiCat by combining an evolutionary scale 300-million parameter ESMC model and a BERT model. The ESMC module represents a parallel model family to ESM3, with a distinct design focus: while ESM3 serves as a generative protein language model, ESMC is optimized for protein sequence representation learning (Hayes *et al*. 2025). Remarkably, ESMC achieves superior performance compared to ESM2 models of equivalent scale, with reduced memory usage, faster inference, and high representational fidelity, making ESMC an excellent choice for feature extraction. In parallel, BERT-based architectures demonstrate the strength of bidirectional Transformer models with attention mechanisms through masked language modeling, enabling them to capture long-range dependencies in protein sequences more effectively (Devlin *et al*. 2019). Further details about the model architecture are described in Materials and Methods.

**Figure 1.**
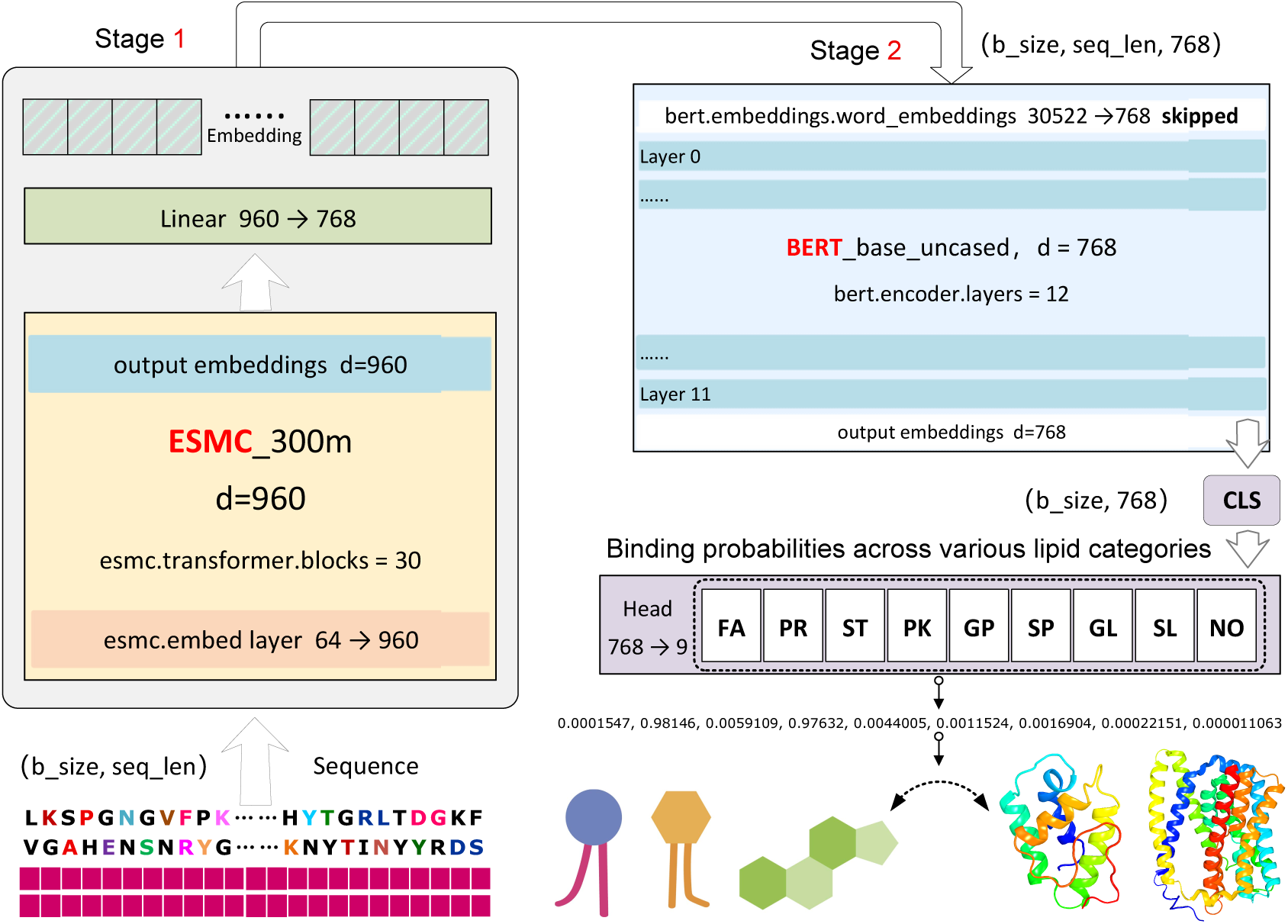
Overall schematic framework of PLiCat. The PLiCat model consists of two stages. In the first stage, the 300M-parameter ESMC_300m model was trained to acquire high-quality embeddings from the input protein sequences. A linear layer (960, 768) was defined for dimensionality reduction. In the second stage, the BERT module takes the output embeddings from the ESMC module to learn relevant protein representations. Finally, the CLS vector from the output of the last Transformer block of the BERT model was fed into a custom classification head to produce a multi-label classification vector. b_size: batch size. seq_len: sequence length.

After training, the [CLS] token embedding, which summarizes the entire sequence, was extracted from the final Transformer block of the BERT model. This embedding was then fed into a custom classification head for sequence-level classification, producing a 9-dimensional vector representing the multi-label predictions. Therefore, PLiCat predicts the probabilities of a given protein sequence binding to any of the eight lipid categories, to multiple classes simultaneously, or to none (**Figure 1**).

The construction of the training and evaluation dataset is based on an open-source protein-lipid database known as BioDolphin (Yang *et al*. 2024). BioDolphin database is a collection of over 127,000 curated protein-lipid interaction entries with corresponding lipid categories (Yang *et al*. 2024). After data cleaning and processing, a dataset comprising 13,673 lipid-binding sequences and 800 non-lipid-binding sequences was utilized for model training. The dataset encompasses a variety of samples as visually depicted in Supplementary Figure 2, allowing for a comprehensive evaluation of lipid-binding patterns under diverse conditions. The organism statistic indicates a diverse species composition, and nearly half of the data come from well-studied model organisms (**Supplementary** Figure 2A). In terms of the distribution of sequence length, the majority of the sequences in the dataset fall within the range of 100 to 400 amino acids, which is consistent with typical protein length distributions observed in natural proteomes (**Supplementary** Figure 2B). Moreover, the lipid-binding proteins can be divided into four different types based on their membrane environment: integral membrane proteins, peripheral membrane proteins, soluble proteins, and lipid-anchored proteins (Ødum *et al*. 2024). The dataset is predominantly composed of soluble and transmembrane proteins, and a smaller proportion of proteins are classified as peripheral or lipid-anchored types, demonstrating the heterogeneity of membrane-association types (**Supplementary** Figure 2C). The diversity of protein data enables the model to learn generalizable features across various species and membrane-association types.

### PLiCat accurately discriminates among lipid-binding categories

In a multi-label classification task, imbalanced data distribution can substantially influence model performance (**Supplementary** Figure 2D). In order to address issues related to an imbalanced dataset as well as avoid overfitting and the long-tail effect, we conducted a 10-fold cross-validation on the training set. Approximately 10% of the dataset was exclusively reserved as an independent test set, and the rest dataset was randomly divided into ten non-overlapping subsets. For each fold, the training and validation sets were divided in a 9:1 ratio (Table S1). Monitoring the training process revealed a consistent trend in different metrics on both training and validation datasets over epochs, suggesting that the model effectively learned stable and generalizable sequence representations without notable degradation in validation performance, thereby supporting the robustness of our training procedure (**Supplementary** Figure 3).

To obtain a rigorous performance evaluation, all ten models trained during the 10-fold cross-validation were tested on the independent dataset representing unseen sequences. Our results reveal that PLiCat achieves superior performance among various metrics, including Accuracy, Precision, Recall, F1 score, Area Under the Curve of the Precision-Recall (AUC-PR), and Area Under the Curve of the Receiver Operating Characteristic (AUC-ROC). All of the performance metrics were consistently high across all folds and with only minor variations observed (**Supplementary** Figure 4). This indicates that the model can generalize well across different subsets of the data with satisfactory robustness and stability.

To obtain the final model for downstream application and evaluation, following the ten-fold cross-validation experiments, we retrained the model from scratch using the entire training dataset, thereby leveraging all available data to maximize learning capacity. To assess model performance under class imbalance, we conducted a detailed assessment across individual lipid categories. The results are summarized in Figure 2C and Table S4. PLiCat achieves excellent performance in AUC-ROC across all categories, with values ranging from 0.88 to 0.97, reflecting high confidence in distinguishing true positives from negatives (**Figure 2A**). Meanwhile, AUC-PR revealed varying performance across different lipid categories, which represents a precision-recall tradeoff, and is more sensitive to class imbalance. Among the eight lipid categories and the negative category, PR (0.91), FA (0.90), and PK (0.91) show the highest precision-recall performance. GP (0.81) and ST (0.83) show moderate performance in AUC-PR curves, while NO (0.74), GL (0.67), SP (0.79), and SL (0.73) displayed relatively lower precision-recall characteristics. Notably, even the lower-performing categories in AUC-PR (e.g., SL, GL) maintained high AUC-ROC values, suggesting the weaker precision-recall tradeoff is likely due to class imbalance (**Figure 2B**).

**Figure 2.**
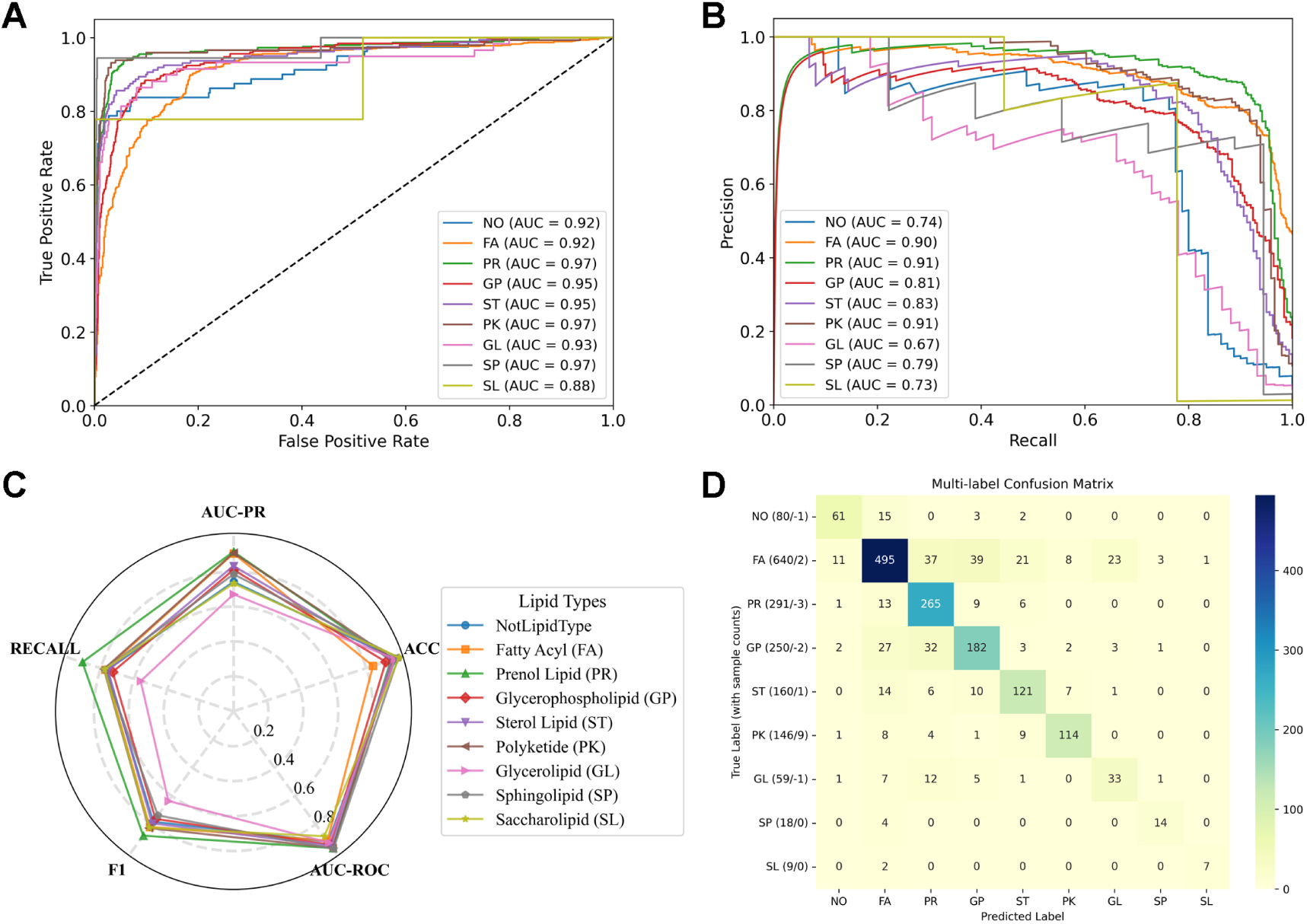
Model performance evaluation. **(A)** Receiver operating characteristic (ROC) curve with area under the curve (AUC) values. **(B)** Precision-recall (PR) curve with area under the curve (AUC) values. **(C)** Radar chart summarizing multiple evaluation metrics. **(D)** Confusion matrix illustrating predictive performance across classes.

The confusion matrix shows correctly predicted samples along the diagonal and misclassified samples on the off-diagonal (**Figure 2D**). The left side of each row indicates the true number of labels, the number of labels the model missed, and the number of labels it overreported. This visualization highlights both the accuracy for each category and the categories that are most frequently confused. Categories such as FA, PR, and PK demonstrate strong performance. In contrast, GP and ST exhibit notable confusion with FA, and GL is occasionally misclassified as PR. Due to the use of a training method based on sample distribution weights, even with a small number of samples (such as SP and SL), still achieved a high accuracy. These results suggest that the model has learned robust features and gained high confidence in the imbalanced dataset.

### Ablation Studies support the effectiveness of ESMC and BERT modules

To validate the effectiveness of different model components, we conducted a series of ablation experiments to demonstrate the contribution of each module component by removing or modifying individual modules while keeping the remaining parts unchanged. Several variant models were devised as follows: First, we removed the pre-trained ESMC module, resulting in the model PLiCat-A. Second, we removed the BERT module, utilizing only the ESMC model for fine-tuning, to generate the model PLiCat-B. Third, we froze all parameters of the pre-trained ESMC module and performed fine-tuning using only the BERT module, termed as PLiCat-C. Fourth, we froze all parameters of the pre-trained ESMC module as well as the first 4 layers of the BERT module, performing fine-tuning with the last 8 layers of the BERT module, termed as PLiCat-D. The final PLiCat-E model was fine-tuned with the last 6 layers of the ESMC module and all layers of the BERT module, while other layers were frozen (**Supplementary** Figure 5).

To examine the impact of different components on model performance, each variant was trained under the same conditions as the PLiCat model. The performance of all the above ablation models was summarized in Table S5. As shown in Supplementary Figure 5G, excluding the ESMC module resulted in a drop in sample accuracy and recall, indicating that ESMC plays a crucial role in capturing lipid-relevant sequence features. Consistently, omitting the BERT encoder and relying solely on the ESMC module led to a less severe effect in performance degradation, suggesting that evolutionary-scale features extracted by ESMC provide complementary information not captured by BERT alone. Similarly, freezing part of the model layers also results in a slight decline in model performance.

The above findings suggest that adjusting the model architecture or the number of frozen layers leads to performance degradation, especially affecting sample accuracy, recall, and F1 score. These results highlight the importance of the individual module for feature extraction, indicating that the designed fusion architecture facilitates the model to efficiently learn the essential lipid-binding signature, and all of the model components play indispensable roles for the model performance.

### PLiCat learned interpretable representations of distinct lipid-binding categories

To better understand how PLiCat captures lipid-binding specificity, we sought to examine the structure of the model’s latent space and assess whether it encodes meaningful, interpretable representations of different lipid-binding categories. First, we analyzed the embeddings of different labels corresponding to single or several lipid categories from the hidden layers using Uniform Manifold Approximation and Projection (UMAP) (McInnes *et al*. 2020). Specifically, we extracted high-dimensional embeddings for each protein sequence via the [CLS] token and projected them into a 2D space. The results reveal that the feature representation of the untrained model is scattered and largely overlapping, while the trained model shows effectively aggregated structures, with clear clustering of sequences into distinct regions (**Figures 3A and 3B**).

**Figure 3.**
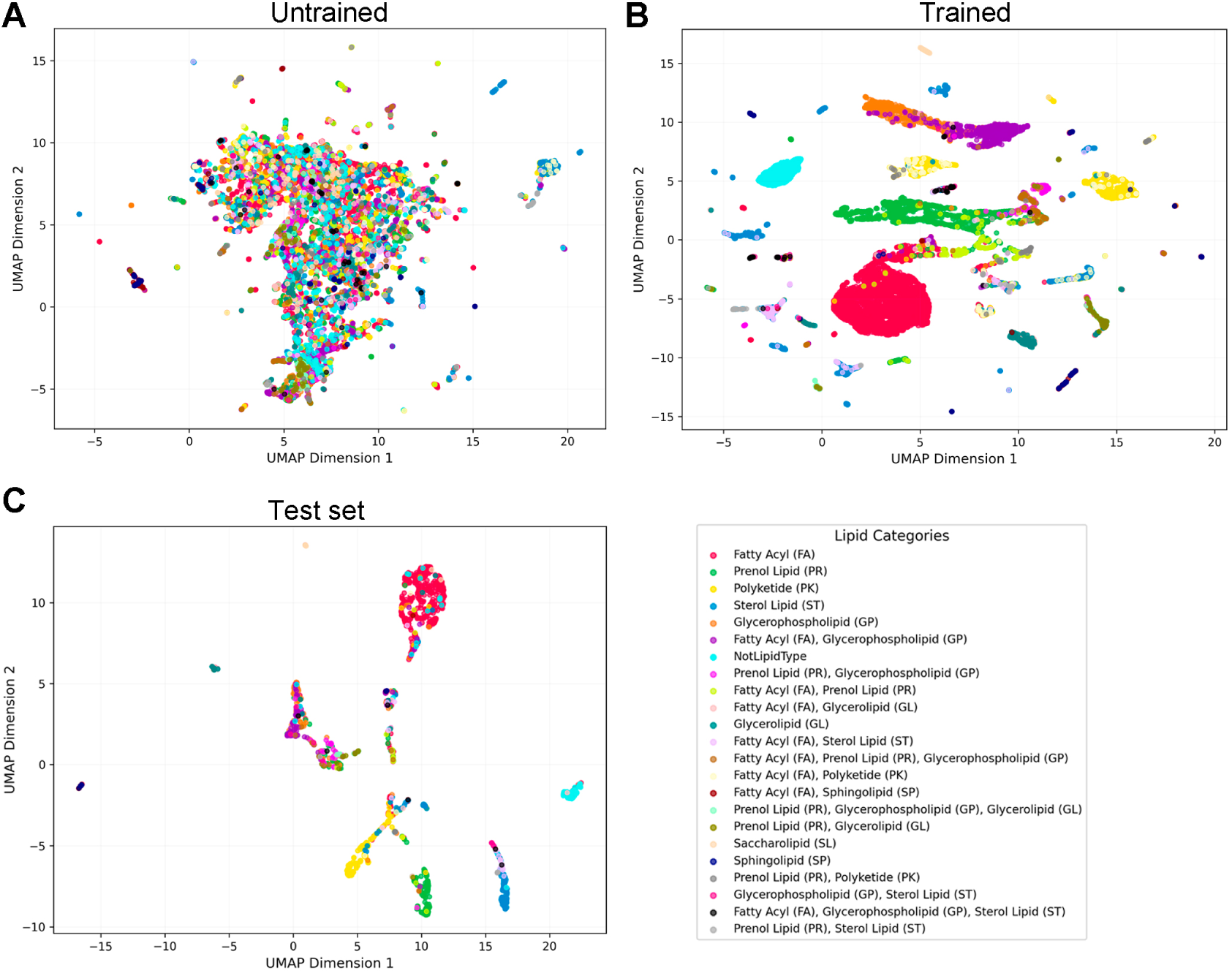
Visualization of the model’s latent space embeddings. **(A)** UMAP projection of sequence embedding distribution before training. **(B)** UMAP projection of sequence embedding distribution after training. **(C)** UMAP projection of sequence embedding distribution for the test set. Each point corresponds to a single data sample, with colors representing different lipid category labels.

Separation between clusters suggests that the model is capable of distinguishing between different classes based on sequence-derived representations. Bridging and overlapping regions may correspond to proteins with shared or ambiguous characteristics (**Supplementary** Figure 6). Furthermore, we analyzed the dimensionality reduction visualization of the embeddings on the test set to assess the model’s generalization. Like in the training set, a similar pattern was observed in the test set, where embeddings form well-separated, compact clusters corresponding to distinct class labels (**Figure 3C**).

To further uncover features that might contribute to selective lipid binding, we analyzed the contribution of individual residues across different lipid categories by computing average attribution scores for each amino acid. Notably, the heat map shows that distinct lipid categories displayed differential binding propensities toward certain amino acid residues. For instance, the amino acid tryptophan (W) shows a positive contribution toward binding with ST and FA, whereas it displays a negative contribution for PR and GP. This pattern illustrates that each lipid category possesses a unique amino acid “signature”, reflecting selective interactions. Some lipid categories, such as SL, showed relatively weak or inconsistent signals across most residues, implying a reliance on global or contextual features rather than specific residue types (**Supplementary** Figure 7). Interestingly, several residues exhibit strong preference regardless of their hydrophobic nature, suggesting that additional factors, such as residue size, charge, polarity, or the local structural context, play important roles in guiding lipid-specific recognition. This highlights the complexity of protein–lipid interactions and suggests that models capturing multi-dimensional residue features can provide deeper insights into lipid selectivity. Together, these results demonstrate that the model’s latent space is not only high-capacity but also interpretable, highlighting the model’s ability to capture meaningful and discriminative features from protein sequences.

### PLiCat facilitates the prediction of lipid-binding residues in two representative proteins

To validate that PLiCat can identify lipid-binding pockets, we applied the Integrated Gradients method to quantify the contribution of each input residue to the model’s prediction. By calculating attribution scores at the residue level, we identified sequence regions that are most influential for the outcome (**Supplementary** Figure 7). We hypothesize that PLiCat’s decision-making relies on sequence features enriched within or near lipid-binding pockets. These high-attribution regions are likely to correspond to residues involved in protein–lipid interactions, thereby allowing indirect inference of lipid-binding sites.

We selected two lipid-binding proteins (UniProt IDs: A9JQL9 and Q6L545) for detailed analysis. As shown in Figures 4A and 4B, residues with top-ranked attribution scores (highlighted with green boxes) partially overlap with the actual binding residues (highlighted with red boxes), indicating that the model can effectively identify lipid-binding sites.

**Figure 4.**
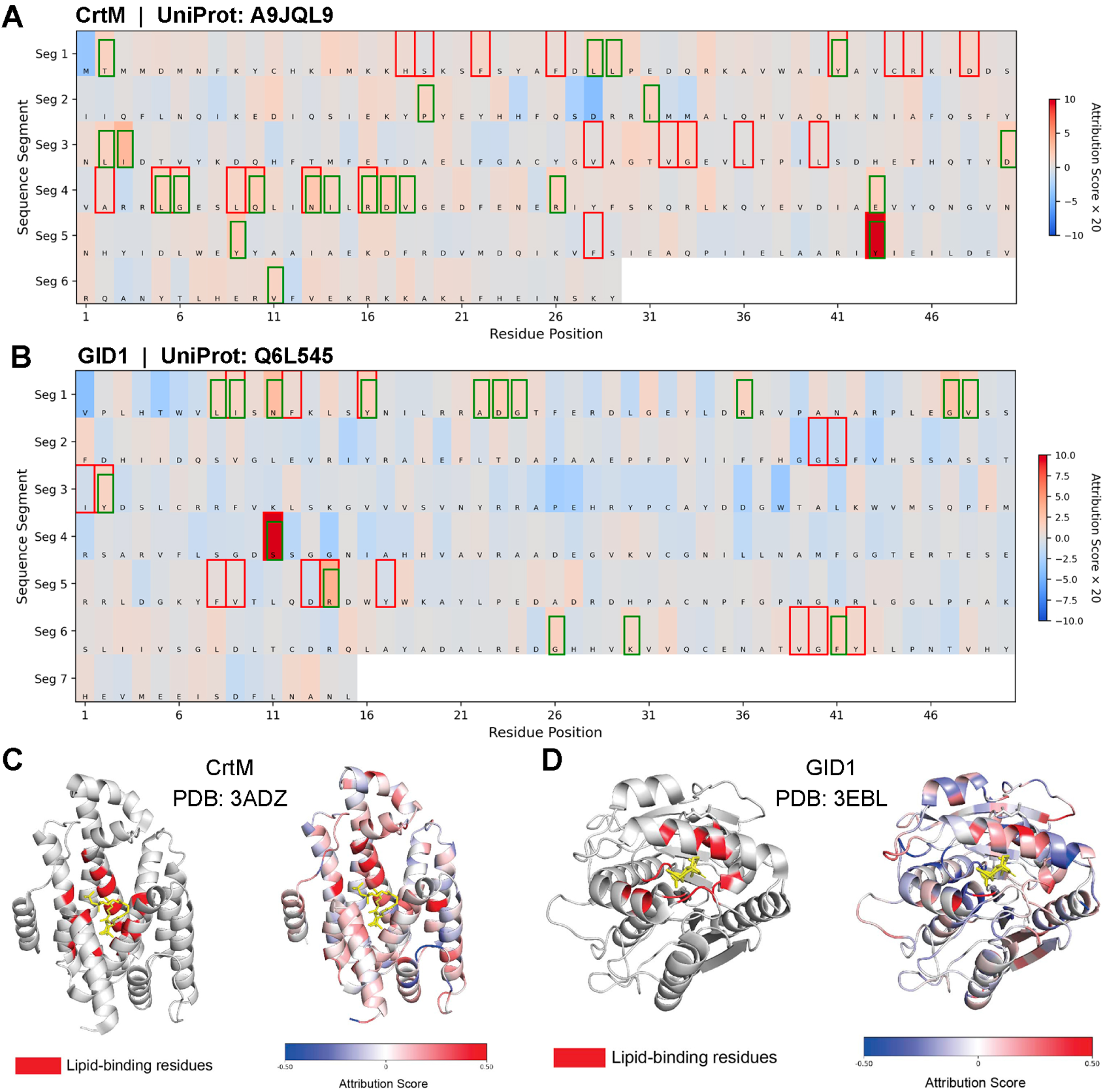
Identification of potential lipid-binding sites by amino acid contributions. **(A, B)** Heatmaps of attribution scores for CrtM and GID1. Heatmaps highlight residues with different attribution scores. Red boxes indicate experimentally validated lipid-binding residues. Green boxes denote the same number of residues with the highest attribution scores outside the red boxes. Overlaps between red and green boxes indicate correctly predicted lipid-binding residues based on the model’s attribution scores. Blue-to-red coloring represents negative to positive attribution scores. **(C, D)** Structural mapping of lipid-binding residues (left) and attribution scores (right) on protein 3D structures. Red regions in the left panels correspond to experimentally determined lipid-binding residues. In the right panels, residues are colored according to their attribution scores (blue: low, red: high). Lipid molecules are shown as yellow sticks. Comparison shows that residues with high attribution scores are spatially enriched around the lipid-binding sites.

A9JQL9, known as Dehydrosqualene synthase (CrtM), is an essential enzyme in *Staphylococcus aureus* that catalyzes the first committed step in staphyloxanthin biosynthesis, which mediates the condensation of two farnesyl diphosphate molecules to form dehydrosqualene (Lin *et al*. 2010, Liu *et al*. 2012, Liu *et al*. 2008). Mapping attribution scores onto the CrtM structures in complex with PSPP, an unreactive substrate analog, demonstrates that residues with higher attribution scores cluster around experimentally determined lipid-binding sites (**Figures 4A and 4C**).

The second protein is GID1 (GIBBERELLIN INSENSITIVE DWARF1), a gibberellin (GA) receptor and a critical component of the GA signaling pathway in plants. GID1 protein specifically bind bioactive gibberellins, triggering conformational changes that facilitate downstream responses (Shimada *et al*. 2008). In the structure of GID1 bound to GA3, the ligand occupies the pocket formed by the N-terminal lid and several loops. Notably, residues contributing most to the model’s prediction closely correspond to the known GA3-binding sites, further supporting the model’s capability to accurately identify true binding sites (**Figures 4B and 4D**). These results suggest the model’s versatility in localizing lipid-binding sites of the input protein sequences, which also provides insights into both the lipid-binding categories recognized by the model and the biological plausibility at the residue level.

### Effects of pathogenic mutations on lipid-binding events

Lipids play fundamental molecular functions in many cellular processes, and dysregulation of lipid metabolism leads to various diseases (Yoon *et al*. 2021). In order to investigate whether lipid-related pathogenic mutations alter lipid-binding or lipid-binding selectivity, we collected a lipid-related pathogenic dataset comprising sequences with mutations. To assess whether wild-type (WT) and mutant (Mut) sequences occupy distinct regions in the model’s representation space, we compared the embeddings and prediction distributions of wild-type and mutant protein sequences. PCA projection of the sequence embeddings reveal that wild-type and mutant samples were largely overlapping in the embedding space, indicating that the model’s internal representations were broadly preserved. A small subset of points is more dispersed in the PCA plot, indicating a possible model-recognized deviation from the canonical sequence behavior (Supplementary Fig. 8A).

To further quantify the shift in prediction profiles, we first computed Jensen– Shannon (JS) divergence over the predicted probability distribution for each lipid category, as a measure of similarity in the model’s predictions between wild-type and mutant sequences. The quantitative divergence analysis reveals label-specific perturbations. Compared to their wild-type counterparts, mutant sequences exhibited marginally higher values for most lipid categories, while SP and SL exhibited minimal changes, indicating the mutations slightly disrupt the model’s prediction in the classification task. Consistently, Wasserstein distance analysis of model logits identified the negative label (NO) and FA as the most mutation-sensitive classes, while other lipid categories exhibited minimal changes. This distributional metric captures the magnitude of change in the model’s output caused by sequence perturbation (**Supplementary** Figures 8B and 8C).

Collectively, these findings suggest that the model captures subtle but meaningful differences introduced by pathogenic mutations. While not all mutations drastically alter predictive outputs, a subset of variants, particularly those with higher Wasserstein distances, appear to disrupt the latent structure of sequence representations in ways that are consistent with functional impact.

## Discussion

The advancement of protein language models has shown tremendous success in a wide range of applications. Protein sequences can be regarded as “sentences” of life which are composed of individual “words” of amino acids. Protein language models can learn the implicit patterns hidden in the “life languages”, therefore enabling prediction and classification. Protein-lipid interactions are universal and crucial for normal cellular activities. Hence, a comprehensive understanding of the protein codes for lipid binding is important for illustrating multiple cellular processes. In this study, we developed a new model to analyze protein-lipid binding, denoted as PLiCat. Leveraging fine-tuned ESMC and BERT models, PLiCat predicts protein-lipid interaction potentials for diverse lipid categories, enabling it to pinpoint sequences critical for lipid-binding selectivity.

Our results indicate that the protein code related to lipid-binding selectivity in natural proteins is likely to serve dual roles: one for correct structural folding and another for binding to specific lipid categories. Over evolutionary timescales, constraints on physicochemical features have likely emerged not only to support protein folding, but also to enable specific lipid-binding.

While PLiCat demonstrates overall strong performance in distinguishing among lipid categories, an examination of Precision-Recall curves reveals disparities in predictive reliability across several lipid categories. Some lipid categories may be underrepresented in the dataset or have similar sequence features, making them harder to distinguish. Therefore, further improvement may require addressing feature overlaps between biologically similar classes and enhancing representation learning for rare categories. These gaps in real-world performance for certain labels (e.g., GL, SL) highlight important considerations for model deployment in an imbalanced context, which merit further investigation and targeted improvement.

There are several key questions pending. First, the interactions between proteins and lipids are not always associated with specific biological functions; in some cases, they may merely serve as structural components within the membrane or complex (Harayama and Riezman 2018, Levental and Lyman 2023). Future studies should focus on elucidating the physiological significance of lipid–protein interactions. Second, our model mainly focuses on eight major lipid categories defined by LIPID MAPS. In addition to major lipid categories, each lipid category can be subdivided into different classes with its subclassification hierarchy (Conroy *et al*. 2024). Our model lacks the higher resolution to discriminate among subclasses within each category, such as selectivity toward lipids with different chain lengths. Improving the ability to resolve such fine-grained differences would be an important direction for future work, as subclass-level specificity is often critical for understanding the biological relevance and providing deeper insights into the molecular mechanisms that govern lipid–protein associations (Harayama and Riezman 2018). For instance, chain length selectivity can directly influence membrane fluidity, signaling pathways, and protein localization, thereby impacting functional outcomes in cellular systems (Levental and Lyman 2023). Third, the selectivity of lipid binding may drive proteins to localize to distinct membrane regions or even alter their intrinsic functions, highlighting the critical role of lipid composition in regulating protein behavior (Levental and Lyman 2023). In the future, leveraging the underlying protein-lipid interaction characteristics learned by the model could be capable to guide the rational design of lipid binders with specific preferences, which represents a valuable research direction.

Taken together, this approach provides an innovative framework and deepens our understanding of protein-lipid interaction. Such efforts could not only serve as a powerful tool for designing proteins with tailored membrane interactions, but also open new avenues for drug development and show promising potential in developing therapeutic strategies for lipid-related diseases.

## Materials and methods

### Datasets

To train the PLiCat model, we collected a lipid-binding protein dataset from BioDolphin. BioDolphin is a curated collection of 127,000 protein-lipid interaction information. We downloaded the BioDolphin dataset from the website (https://biodolphin.chemistry.gatech.edu/). For dataset construction, we extract lipid-binding protein sequences and the corresponding lipid categories. The sequences in BioDolphin possess a considerable degree of redundancy. To generate a high-quality dataset, we performed group-by and removed duplicates based on protein sequences. Furthermore, sequences containing illegal characters were replaced by the canonical sequence obtained from UniProt. After this data cleaning process, we collected a lipid-binding protein dataset, resulting in 16112 proteins with lengths between 35 and 500 amino acids. Given the considerable imbalance between different lipid categories, we analyzed the distribution of samples across different labels in the lipid-binding protein dataset. Labels occurring less than 50 times were filtered out, and their corresponding samples were excluded from the dataset, resulting in a final positive lipid-binding protein dataset containing 13673 samples.

For negative samples, the non-lipid-binding proteins were collected from UniProt using stringent criteria, yielding 7150 negative samples with sequence length between 35 and 500 amino acids. To reduce redundancy, negative samples were clustered using the CD-HIT algorithm with an identity threshold of 70% (Li and Godzik 2006). Subsequently, a total of 800 sequences were randomly selected from the filtered negative samples to maintain relative balance across the dataset.

The final dataset contains 12873 positive samples and 800 negative samples. To ensure clear separation between classes, we confirmed that the positive and negative samples are mutually exclusive. The statistical information of the dataset is shown in Supplementary Figure 2.

For data splitting, 10% of the proteins were selected to produce the independent test dataset, which was used solely for testing. The remaining 90% dataset was used for 10-fold cross-validation. The final training and testing datasets contain 12296 and 1377 sequences, respectively (Table S1).

### Model architecture, training, and evaluation

The PLiCat model consists of two major components: ESMC and BERT. The pre-trained ESMC model contains approximately 300 million parameters and produces an embedding dimension of d_ESMC_=960. The ESMC model comprises 30 Transformer blocks that employ Rotary Position Embedding (RoPE) for efficient encoding of positional information. Within each Transformer block, the feed-forward network (FFN) undergoes a dimensionality transformation of (960,5120)→(2560,960). The shape of the input embedding layer is (64,960), where 64 corresponds to the amino acid vocabulary size defined by ESM. Next, the representation from ESMC was then passed to the BERT model. Due to the embedding dimension of BERT is 768, a linear transformation of size (960, 768) was defined to reduce the dimensionality. We fine-tuned all the transformer layers of the encoder of ESMC and BERT.

We employed a two-stage training and evaluation strategy consisting of 10-fold cross-validation followed by final model selection and evaluation on a held-out test set. For model training, PLiCat was trained end-to-end with a batch size of 16 and evaluated using weighted binary cross-entropy loss with logits (BCEWithLogitsLoss). Optimization was performed using the AdamW algorithm with default parameters and an initial learning rate of 2e-5. More details about the model and training parameters are summarized in Table S2 and Table S3. All models were implemented in PyTorch 2.7.

To address the imbalance between positive and negative samples in the training set, we adopted a weighted loss function that assigns higher weights to rare classes, thereby increasing their contribution to the training objective. The class-specific weight was calculated according to the following equation:

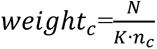

Where *weight_c_* denotes the weight assigned to class c, N is the total number of samples in the training set, K is the total number of classes, and n_c_ is the number of positive samples in class c.

Using a weighted loss function during training is appropriate for mitigating class imbalance; however, to objectively evaluate the model’s generalization performance, we used the unweighted loss and metrics on the validation set. This avoids artificially amplifying the influence of rare classes during evaluation. For model evaluation, the model performance was assessed using multiple metrics, including AUC-ROC, AUC-PR, F1 score, Accuracy, and Recall. Accuracy was computed at both the label level and sample level. We reported macro-averaged scores across labels unless otherwise specified. For each fold, the model was trained on 90% of the fold data and validated on the remaining 10%. This procedure produced ten independently trained models, which were subsequently used to perform inference on the test dataset, enabling assessment of performance robustness and variability across folds.

### Calculation of attribution scores

To interpret the contribution of each residue in the input protein sequence to the model’s prediction, we employed Integrated Gradients (IG) as implemented in Captum (v0.8.0). IG computes the integral of gradients with respect to a baseline input, attributing importance scores to each amino acid based on its contribution to the model output. We used an all-unknow (<UNK> token) sequence as the baseline input and approximated the path integral using 50 steps.

### Latent space investigation of PLiCat

To investigate the structure of learned representations, we extracted latent embeddings of protein sequences from the trained model. Latent space analysis was performed on both training and test sets. For each protein, we extracted its latent embeddings from the final encoder layer. To visualize the structure of these high-dimensional embeddings, the Uniform Manifold Approximation and Projection (UMAP) method is implemented using the umap-learn library (0.5.9.post2) with default hyperparameters. In each UMAP projection, individual protein is shown as point and colored by their lipid-binding categories annotation.

### Pathogenic variants analysis

We developed a lipid-related pathogenic protein dataset specifically tailored to analyze the pathogenic effects. The pathogenic mutations dataset was sourced from multiple databases. Variants linked to Mendelian diseases were obtained from the ClinVar Variant Database (Landrum et al. 2018). Various Cancer-associated studies were available through cBioPortal, including The Cancer Genome Atlas (TCGA) and Therapeutically Applicable Research to Generate Effective Treatments (TARGET) (Cerami et al. 2012, Consortium et al. 2017). Genomic coordinates of cancer variants originally based on the hg19 reference genome were converted to hg38 using Liftover (Kent et al. 2002). To restrict our focus to lipid-related diseases, Ensembl VEP v114 was used for variant annotation (Harrison et al. 2020). Synonymous mutations, duplication mutations, intron/UTR/regulatory region mutations were excluded, and DNA variants that led to the same amino acid change were treated as equivalent.

### Computation of JS Divergence and Wasserstein Distance

Jensen–Shannon divergence and Wasserstein distance were computed to quantify the differences between model outputs for wild-type and mutant protein sequences. For the Jensen–Shannon divergence, the predicted probability distributions P (wild-type) and Q (mutant) for each label were first obtained from the model outputs. The divergence was calculated as:

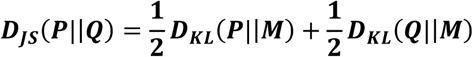

Where 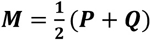 is a mixture distribution of P and Q.

The Wasserstein distance (Earth Mover’s Distance) was calculated to measure the minimal “cost” of transforming one distribution into another. For each label, the model’s unnormalized outputs (logits) for the wild-type and mutant sets were treated as empirical distributions. The first-order Wasserstein distance *W (P, Q)* was computed as:

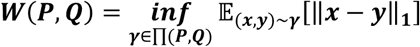

Where P, Q are two distributions, ∏(***P, Q***) denotes the set of all possible joint distributions with marginals P and Q. ‖*x-y*‖ represents the transportation cost on the real number line.

### PCA projection analysis

Principal Component Analysis (PCA) was applied to visualize and compare the distribution of model embeddings derived from wild-type and mutant protein sequences. For each sequence, the final-layer embedding vector from the trained model was extracted and standardized to have zero mean and unit variance across each feature dimension. PCA was performed using the scikit-learn implementation, with the first two principal components (PC1 and PC2) used for two-dimensional visualization. The total explained variance was calculated as the cumulative proportion of variance captured by the selected components, reflecting the extent to which the original feature variance was preserved in the reduced-dimensional space. The scatter plots of PC1 versus PC2 were generated to examine clustering patterns and potential distributional shifts between wild-type and mutant embeddings.

## Data availability

The datasets used for training and testing are available at https://github.com/Noora68/PLiCat/tree/main/process_data.

## Code availability

The model source code is available at https://github.com/Noora68/PLiCat. The trained PLiCat model is available at https://huggingface.co/Noora68/PLiCat-0.4B.

## Acknowledgements

We would like to appreciate Professor Maofu Liao for the insightful advice and careful review on this work, which greatly improved the quality of this paper. We also acknowledge the use of computing resources located at Ludong University. Last but not least, thanks to all open-source programming libraries and databases.

## Author Contributions

F.D. conceived the project. F.D. designed and performed the computational analyses. F.D. wrote the manuscript. J.W. contributed to manuscript review and revision. All authors collaboratively discussed the results, and agreed on the content of the paper.

## Conflict-of-interests

The authors declare no competing interests.

## Supplementary Figures and Tables

**Supplementary Figure 1.**
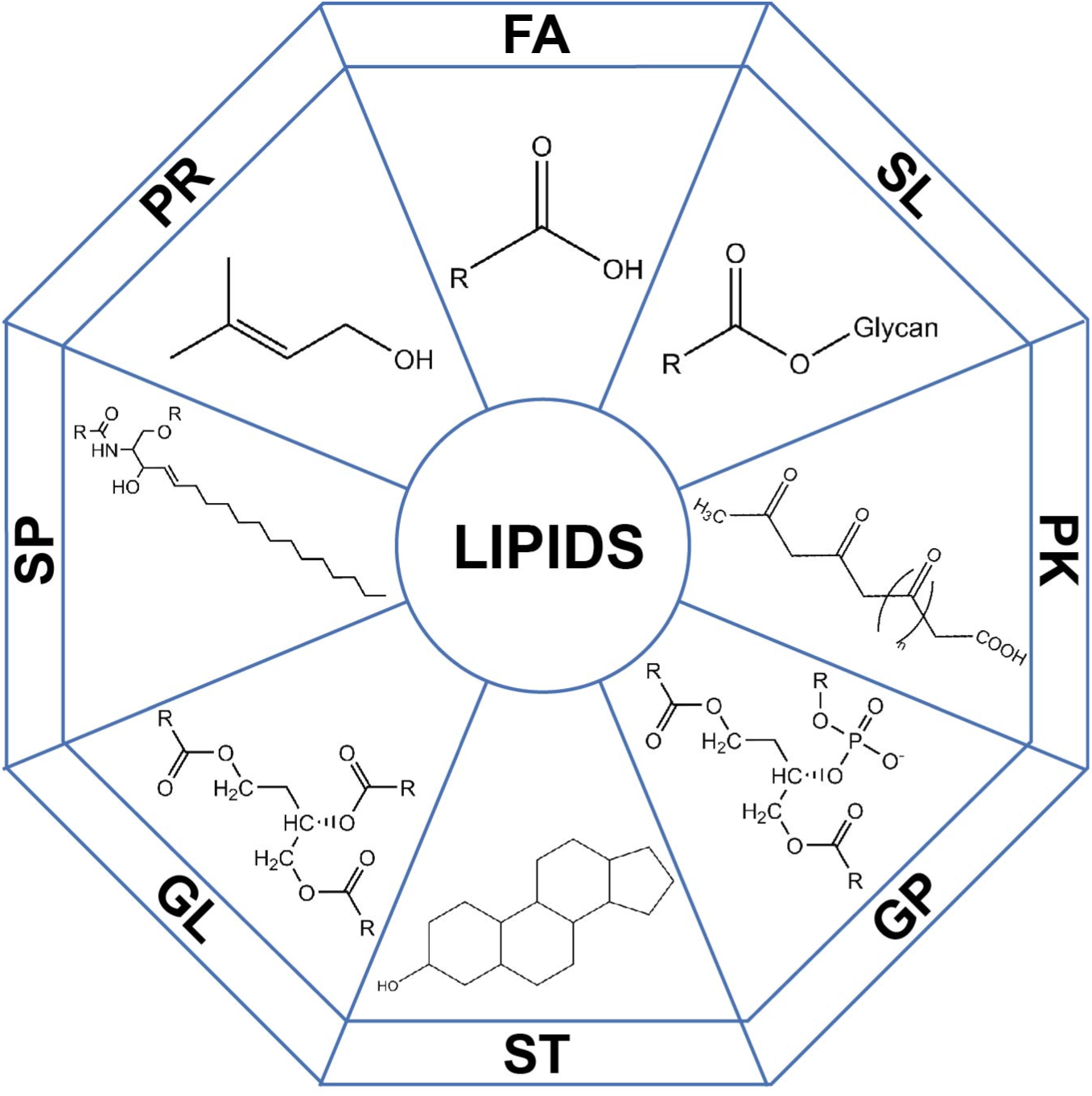
Lipid classifications. Graphical depiction of different lipid categories defined by LIPID MAPS. Fatty Acyls (FA), Glycerolipids (GL), Glycerophospholipids (GP), Sphingolipids (SP), Sterol lipids (ST), Prenol lipids (PR), Saccharolipids (SL), and Polyketides (PK).

**Supplementary Figure 2.**
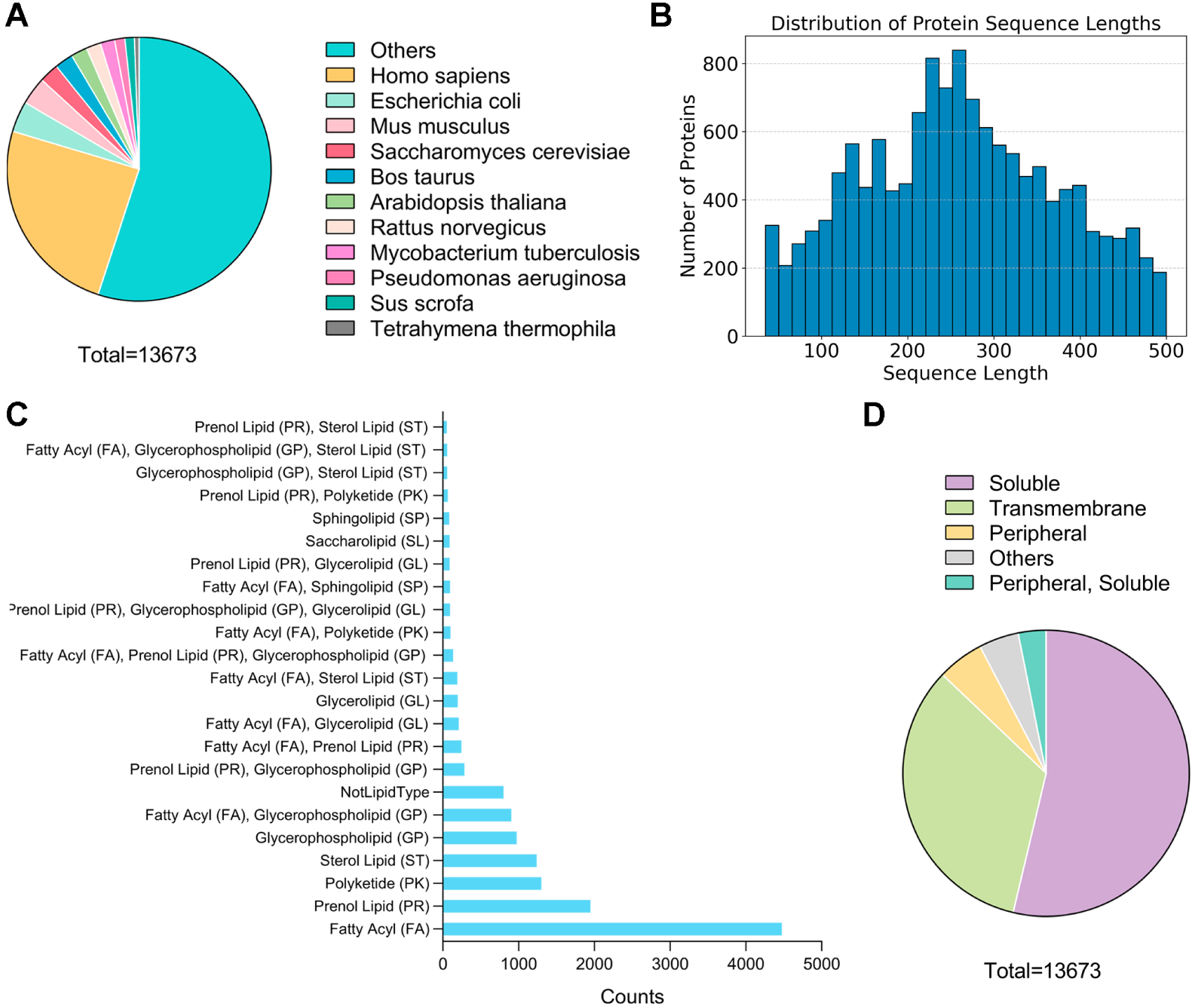
Data distribution. **(A)** Source organism distribution of the dataset, where major model species are shown individually and all other species are grouped under “Others”. **(B)** Statistical distribution of sequence lengths within the dataset. **(C)** Distribution of lipid-binding category labels. **(D)** Distribution of membrane-association types. Annotation of membrane association types was obtained from UniProt and supplemented with DeepLoc 2.1 (Ødum et al. 2024).

**Supplementary Figure 3.**
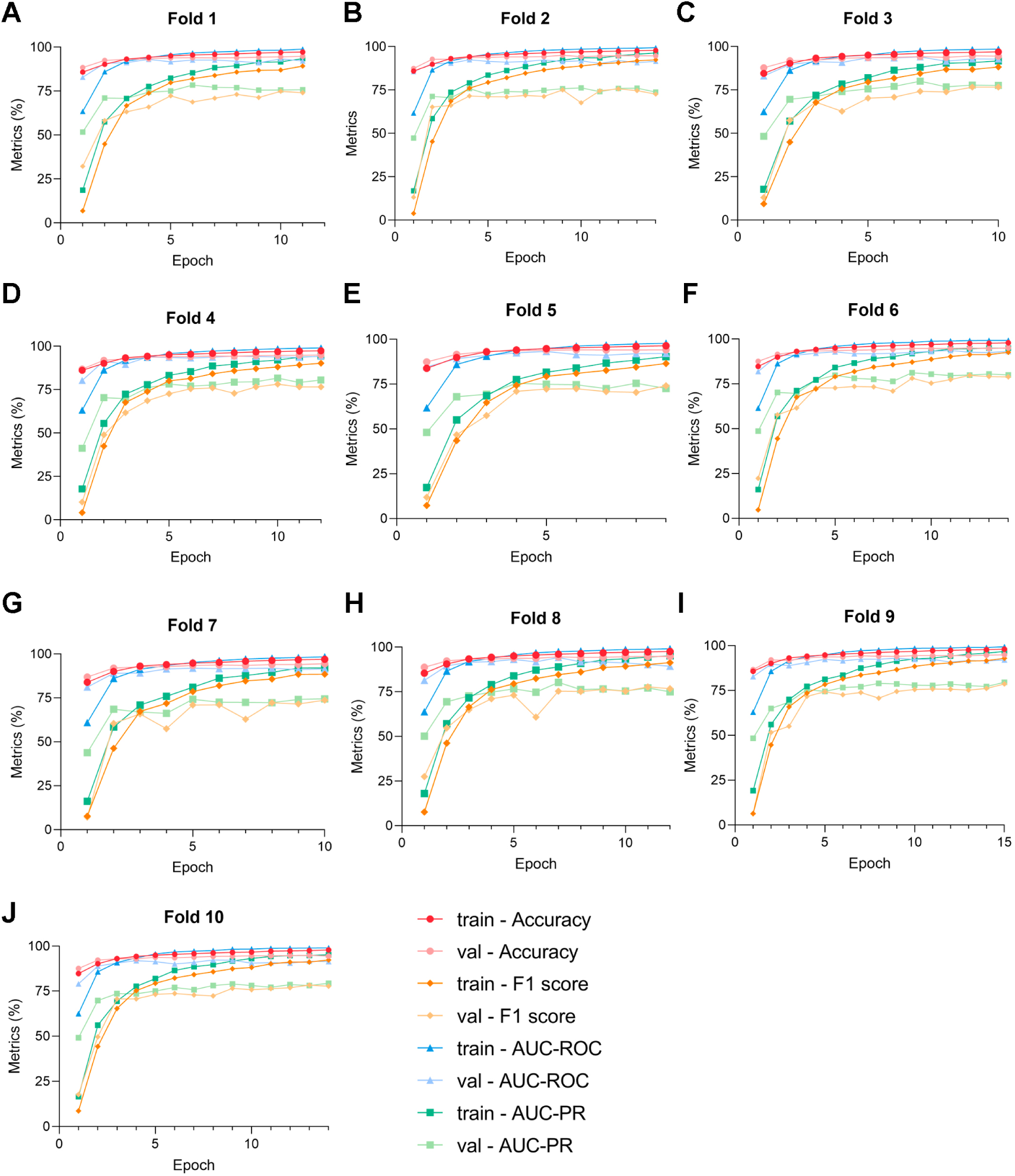
Monitoring of the metric trajectories of the 10-fold cross-validation. For each fold, Accuracy, F1 score, AUC-ROC, and AUC-PR were recorded after each training epoch, illustrating the evolution of model performance over the training process. (A-J): Fold 1 – Fold 10. val: validation set.

**Supplementary Figure 4.**
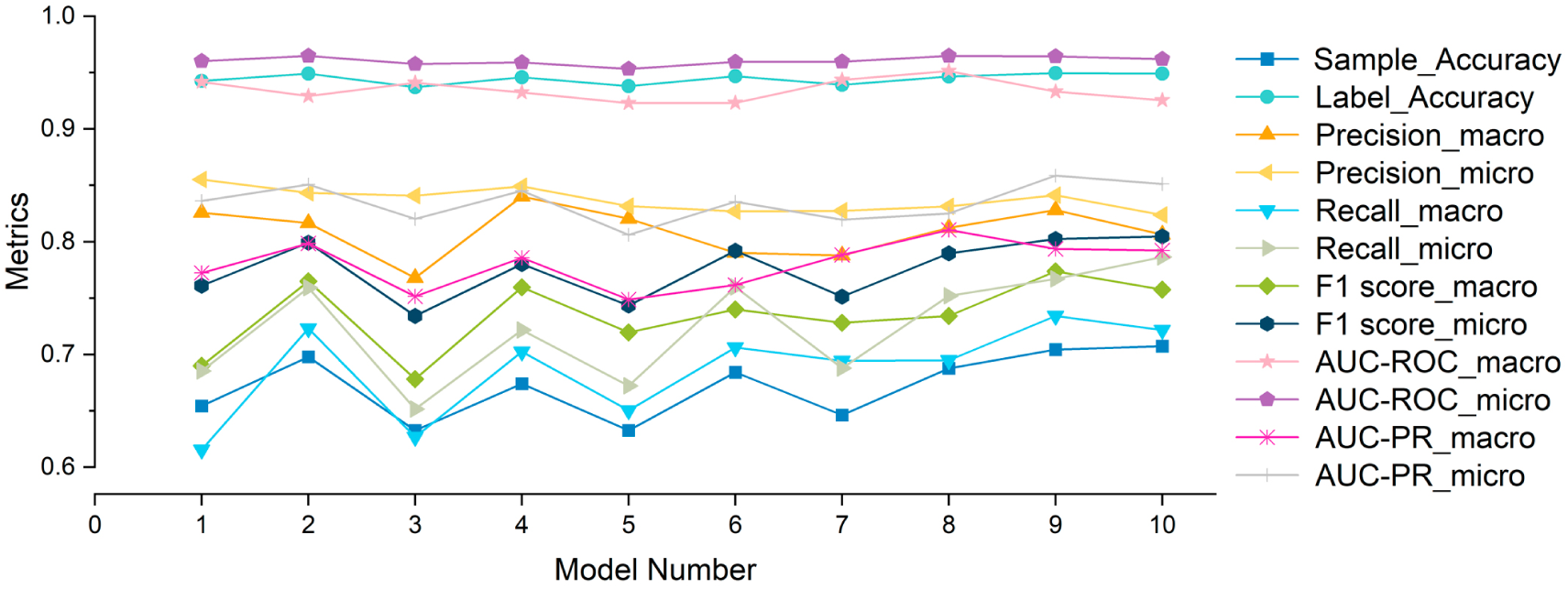
Comparison of evaluation metrics across models obtained from 10-fold cross-validation. Each point represents the evaluation of one of the ten models trained during 10-fold cross-validation. Metrics are shown for both micro-averaged and macro-averaged calculations, including Accuracy (Sample and Label), Precision, Recall, F1 score, Area Under the Curve of the Receiver Operating Characteristic (AUC-ROC), and Area Under the Curve of the Precision-Recall (AUC-PR).

**Supplementary Figure 5.**
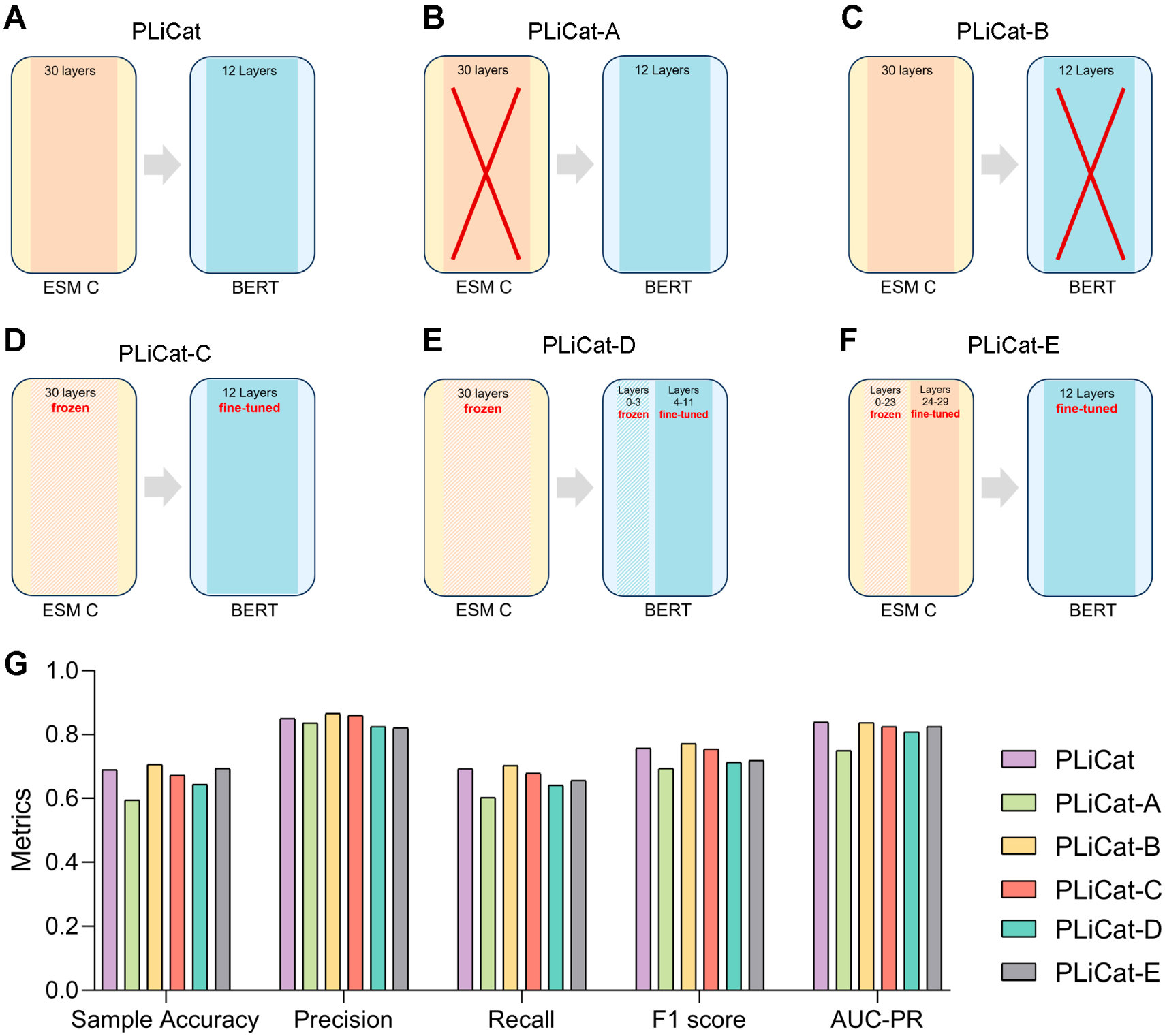
Ablation Study. (A-F) Schematic diagrams of different ablation models. **(A)** The original PLiCat model architecture. **(B)** PLiCat-A: Removing the ESMC module, fine-tuning using only the BERT module. **(C)** PLiCat-B: Removing the BERT module, fine-tuning using only the ESMC module. **(D)** PLiCat-C: Using a pre-trained frozen ESMC model and fine-tuning the BERT model. **(E)** PLiCat-D: Freezing the ESMC module and the firt 0-3 layers of the BERT module, fine-tuning the last 8 layers of the BERT module. **(F)** PLiCat-E: Freezing the first 24 layers of ESMC, fine-tuning the last 6 layers of ESMC, and the BERT module. **(G)** Comparative performance of different ablation models across multiple evaluation metrics.

**Supplementary Figure 6.**
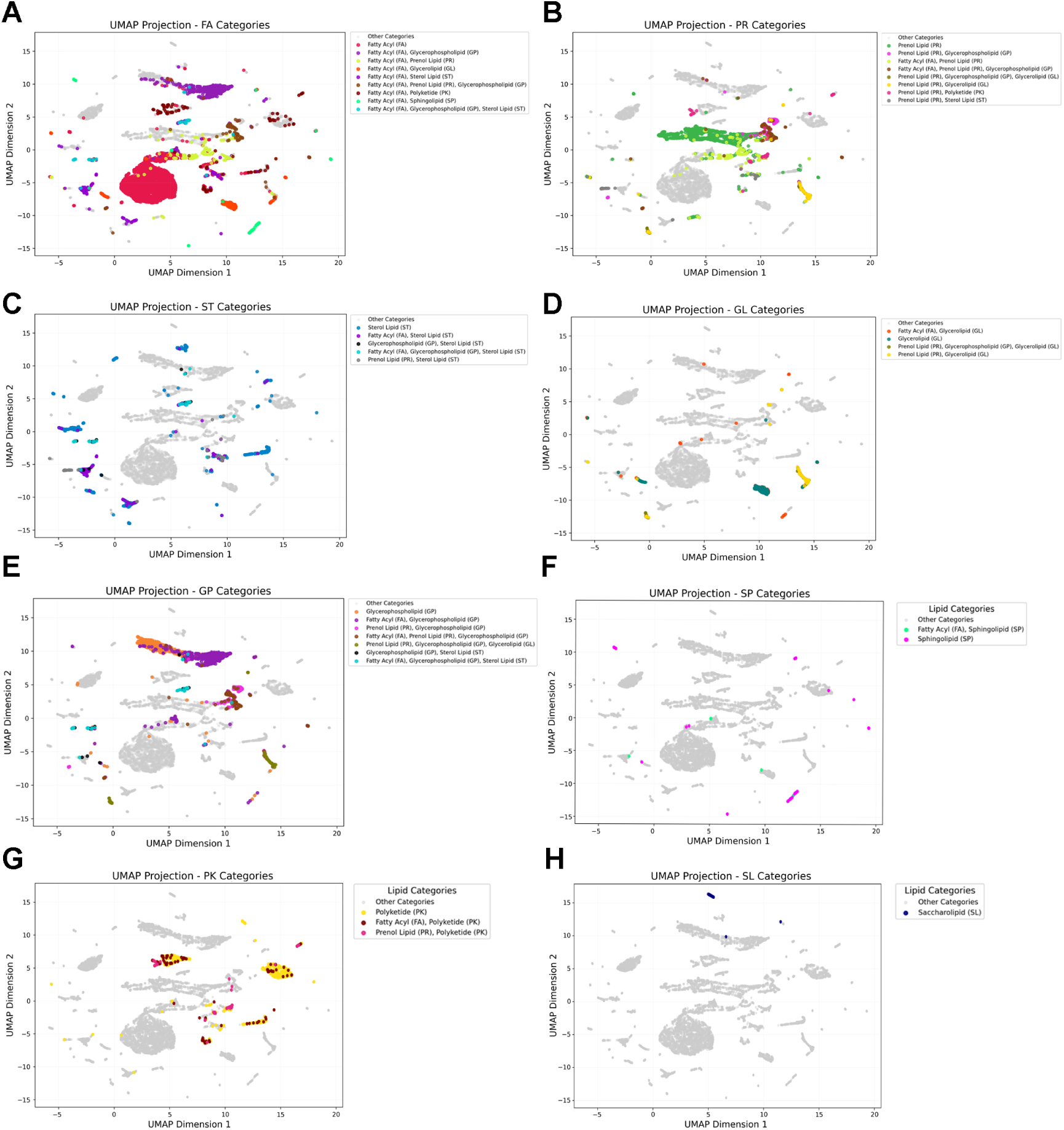
UMAP projection of protein embeddings for different lipid categories. Each subpanel (A-H) highlights a specific lipid category (colored dots) while other categories are shown in gray. **(A)** Fatty Acyl (FA, red), **(B)** Prenol Lipid (PR, green), **(C)** Sterol Lipid (ST, blue), **(D)** Glycerolipid (GL, cyan), **(E)** Glycerophospholipid (GP, orange), **(F)** Sphingolipid (SP, purple), **(G)** Polyketide (PK, yellow), and **(H)** Saccharolipid (SL, navy).

**Supplementary Figure 7.**
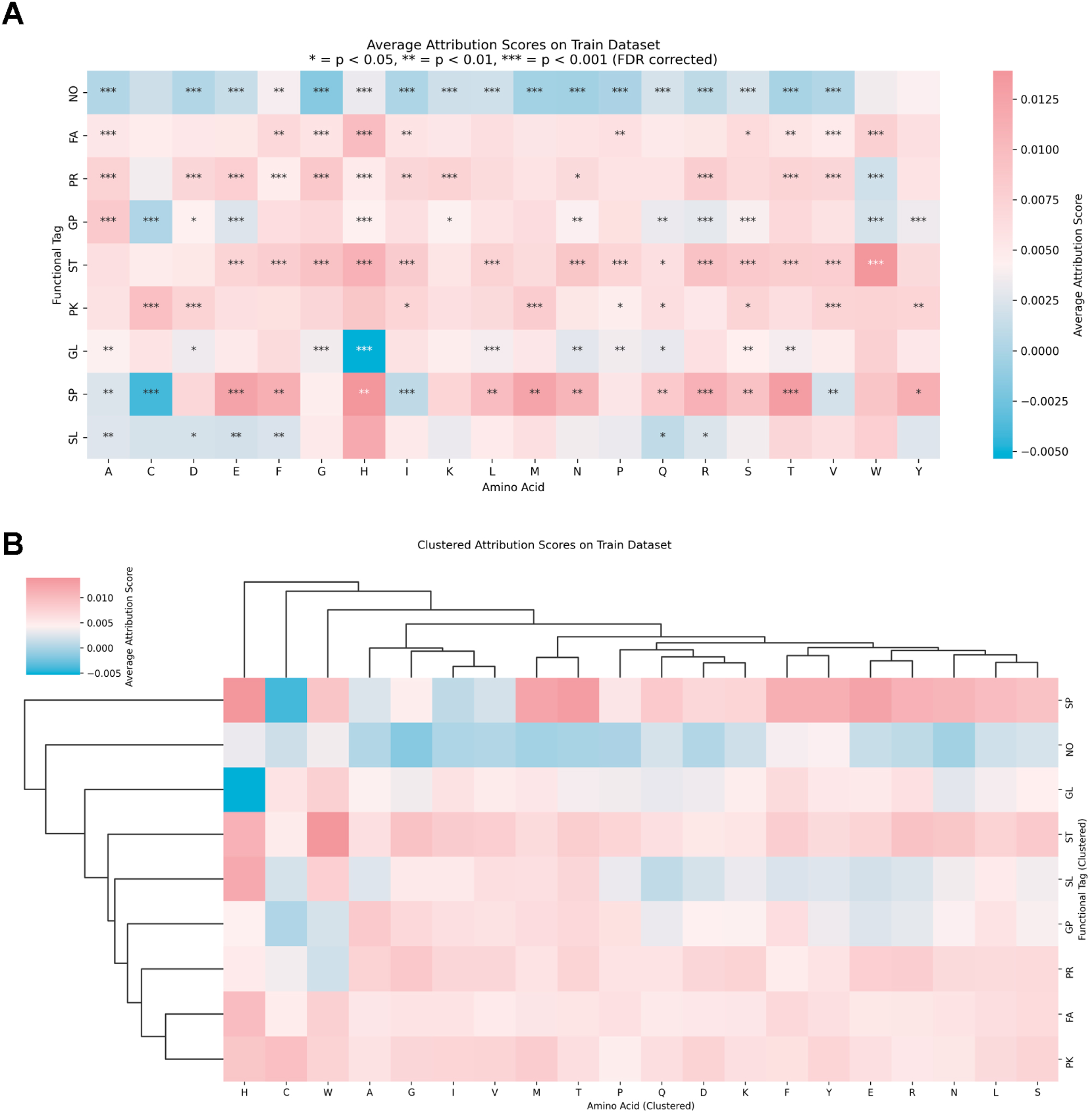
Attribution scores of amino acids for different lipid-binding categories predictions. **(A)** Heatmap visualization of attribution scores, illustrating the contribution of each amino acid to the prediction for different lipid categories. Rows correspond to amino acids. Columns represent 8 lipid categories and negative category. **(B)** Clustering of different categories by attribution scores of amino acids.

**Supplementary Figure 8.**
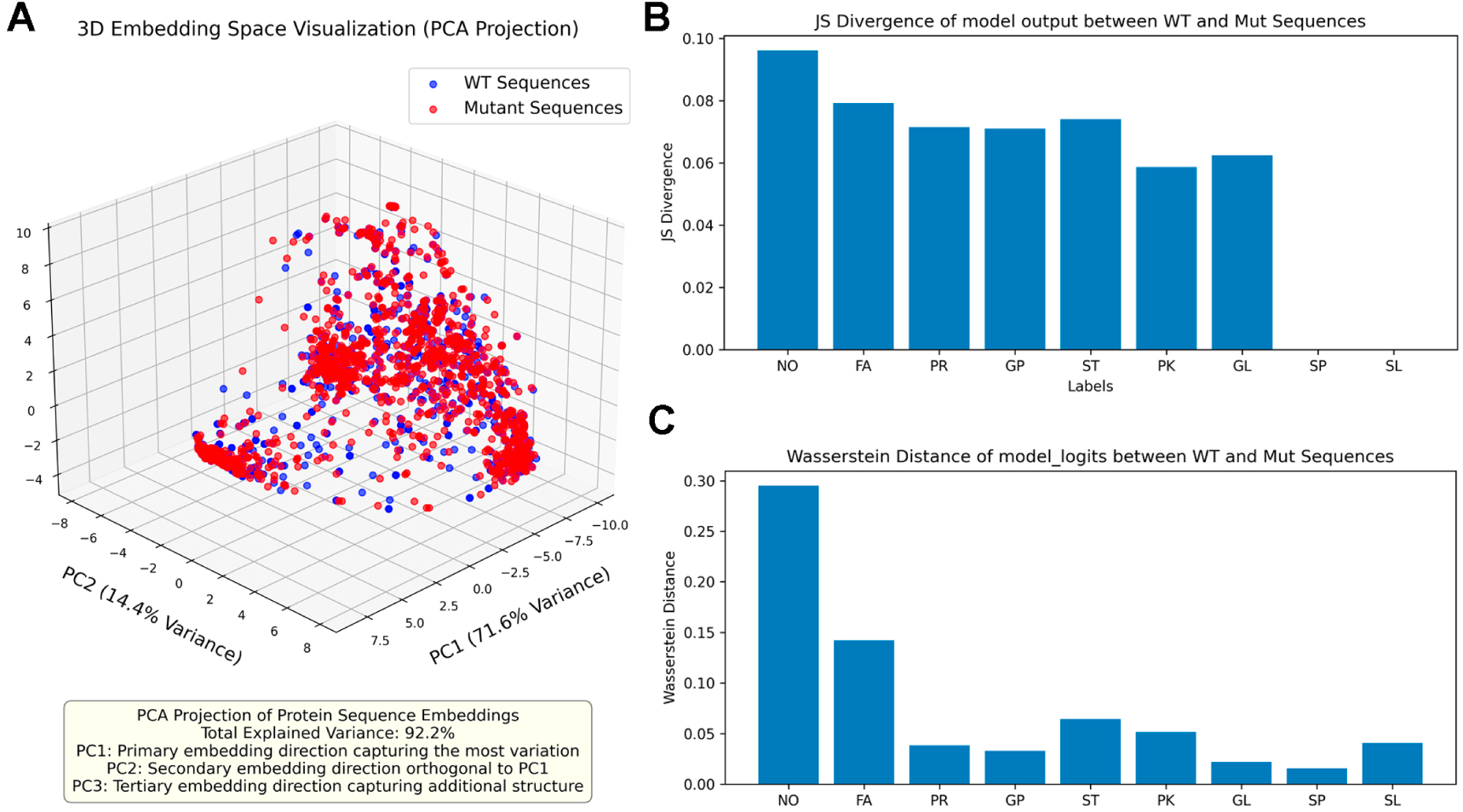
Effects of pathogenic mutations on lipid binding events. **(A)** PCA projection of the embeddings from wild-type (WT) and mutant (Mutant) sequences. **(B)** Jensen–Shannon (JS) divergence computed between wild-type (WT) and mutant (Mutant) datasets for each category. **(C)** Wasserstein distance computed between wild-type (WT) and mutant (Mutant) datasets for each category.

**Table S1.** Positive and negative data for training and testing.

| Dataset | Number of data |  |
| --- | --- | --- |
|  | Positive | Negative |
| Training set | 11576 | 720 |
| Test set | 1297 | 80 |

See methods for more detailed information.

**Table S2.** Architectural Hyperparameters of PLiCat.

| Module | Parameter | Value | Description |
| --- | --- | --- | --- |
| ESMC | Number of Transformer Blocks | 30 | Model depth |
|  | Hidden layer vector dimension | 960 | Model width |
|  | Input embedding dimension | (64, 960) | 64 corresponds to vocabulary size |
|  | ESMC FFN | (960, 5120)<br>→ (2560, 960) | Asymmetric two-layer linear module |
|  | Number of attention heads | 15 | Enhance the model's representational capacity |
|  | Per-head dimension | 64 | Vector dimension divided by the number of attention heads |
|  | Activation function | SwiGLU | Improve training stability |
|  | Linear | (960, 768) | From ESMC to BERT |
| BERT | Number of BERT-Base Uncased Transformer Blocks | 12 | Model depth |
|  | Hidden layer vector dimension | 768 | Model width |
|  | Input embedding dimension | (30522, 768) | Replaced by dimensionally reduced (960, 768) |
|  | Number of attention heads | 12 | Enhance the model's representational capacity |
|  | Per-head dimension | 64 | Vector dimension divided by the number of attention heads |
| | FFN | (768, 3072)<br>→ (3072, 768) | Dimension: $d \rightarrow 4d \rightarrow d$ |
|  | Activation function | GELU | Continuously differentiable, avoiding the gradient discontinuity present in ReLU |
|  | BERT classification head | (768, 9) | Multi-label classification vector |
| Number of Parameters | Total parameters | 443,818,057 | Relate to model architecture |
|  | Trainable parameters | 443,818,057 | Relate to fine-tuning strategy |

**Table S3.** Training Hyperparameters of PLiCat.

| Parameter | Value | Description |
| --- | --- | --- |
| Training strategy | 10-fold cross-validation | Evaluate the model's generalization ability |
| Dataset split | Proportional to the label distribution | Preserve class distribution in subsets |
| Optimization algorithm | AdamW | $\beta_1 = 0.9, \beta_2 = 0.95$ |
| Loss Function | BCEWithLogitsLoss | Weighted computation of training loss |
| Weight decay | 0.05 | L2 regularization strength |
| Maximum sequence length | 500 | Context window |
| Early stopping strategy | Validation loss is not decreasing for five consecutive epochs | Identify the optimal model and avoid overfitting |
| Learning rate scheduling strategy | initial learning rate: $2e-5$, with a warmup + cosine decay schedule | Warmup phase accounts for 10% of the total training steps to improve stability in the early stages, followed by a cosine decay phase over the remaining 90% to enhance generalization in the later stages |
| Gradient clipping | Maximum gradient norm of 1.0 | Improve training stability |
| Threshold for multi-label binary classification | 0.6 | Sigmoid function to compute the logits of the model outputs |

**Table S4.** PLiCat performance metrics of different lipid categories.

|  | Accuracy | Precision | Recall | F1 score | AUC-ROC | AUC-PR |
| --- | --- | --- | --- | --- | --- | --- |
| FA | 0.840232 | 0.868421 | 0.773438 | 0.818182 | 0.91558 | 0.904602 |
| PR | 0.947712 | 0.85209 | 0.910653 | 0.880399 | 0.968743 | 0.914436 |
| GP | 0.917211 | 0.798246 | 0.728 | 0.761506 | 0.945132 | 0.811365 |
| ST | 0.954248 | 0.834483 | 0.75625 | 0.793443 | 0.952558 | 0.833522 |
| PK | 0.965142 | 0.876923 | 0.780822 | 0.826087 | 0.967256 | 0.907716 |
| GL | 0.972404 | 0.733333 | 0.559322 | 0.634615 | 0.930429 | 0.670354 |
| SP | 0.992738 | 0.7 | 0.777778 | 0.736842 | 0.973878 | 0.785799 |
| SL | 0.997821 | 0.875 | 0.777778 | 0.823529 | 0.884747 | 0.732134 |
| NO | 0.975309 | 0.802632 | 0.7625 | 0.782051 | 0.923174 | 0.742193 |

**Table S5.**
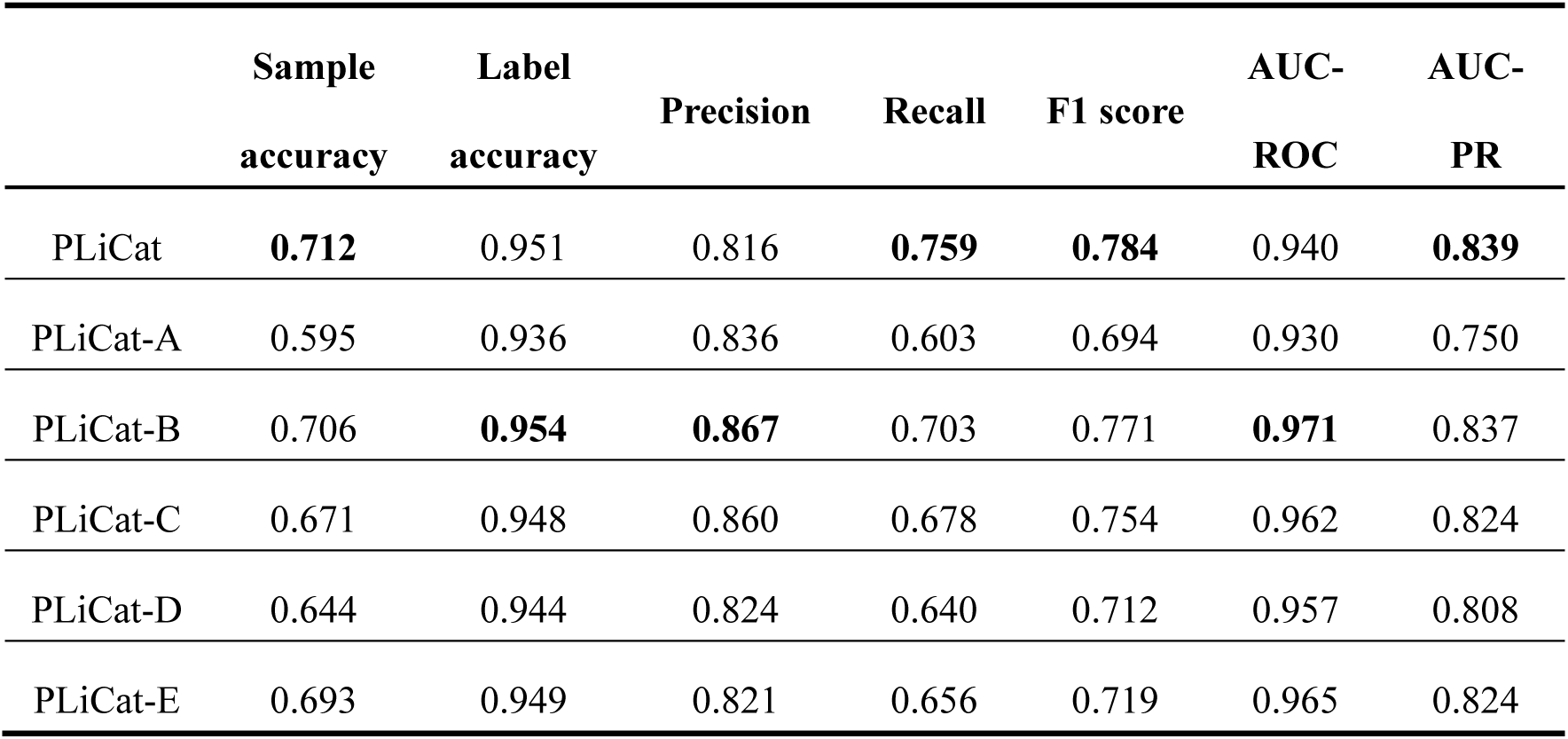
Ablation Studies.

## Notes

https://github.com/Noora68/PLiCat

https://huggingface.co/Noora68/PLiCat-0.4B

### Competing Interest Statement

The authors have declared no competing interest.

